# Essential Role of B Cells in Early Defense Against *Acinetobacter baumannii* Pulmonary Infections

**DOI:** 10.1101/2025.01.24.634778

**Authors:** Aminul Islam, Luis A. Actis, Timothy J. Wilson

## Abstract

Treatment of antibiotic-resistant *Acinetobacter baumannii* infections has become exceedingly challenging, leading to higher morbidity and mortality. Therefore, the development of new therapeutics to treat these infections is critically needed, and immunotherapy is one potential treatment option. Immune responses against this pathogen, particularly lymphocyte-mediated responses, are not well understood. In this study, we investigated the role of B cells in innate resistance to *A. baumannii* pulmonary infection using a B cell-deficient (µMT) mouse model. B cell-deficient mice were impaired in clearing A. baumannii from the lung, liver, and spleen and failed to prevent extrapulmonary dissemination of *A. baumannii* after infection. Transcriptomic analyses indicated that the lack of B cells was associated with reduced expression of several genes encoding antimicrobial proteins. In addition, B cell deficiency was associated with pulmonary eosinophilia and increased pulmonary recruitment of Ly6C^+^ NK cells following *A. baumannii* infection. This study demonstrates the significant role of B cells in providing early protection against *A. baumannii* infection and implicates B cell-dependent mechanisms beyond antibody-mediated bacterial resistance.

## Introduction

*Acinetobacter baumannii* is a Gram-negative coccobacillus that has emerged as one of the most serious public health threats globally. Although this nosocomial pathogen is associated with diverse types of infection, including meningitis, bloodstream, wound, and urinary tract infections, ventilator-associated pneumonia is the most severe form of infection, with a mortality rate of up to 60% (1). Because of the increasing emergence of pan-drug-resistant strains, treatment of *A. baumannii* infections has become very challenging, leading to high morbidity and mortality (2, 3). Drug-resistant *A. baumannii* was associated with approximately 500,000 deaths in 2019 (4). Moreover, multidrug-resistant *A. baumannii* is often associated with serious secondary infections in hospital settings. For example, COVID-19 patients with secondary *A. baumannii* infections experienced a two-fold higher mortality rate (5). Because of the increased drug resistance and severity of infections, the World Health Organization categorized carbapenem-resistant *A. baumannii* as a priority one threat to human health for which new therapeutic agents need to be developed immediately (6). Understanding A. baumannii-host interactions, including immune responses against this opportunistic pathogen, is essential for developing effective therapeutics.

Studies have demonstrated the important protective role of innate immune effectors against *A. baumannii* (7). Among innate immune cells, neutrophils play a dominant role in clearing *A. baumannii* infection. Neutrophil deficiency leads to increased lethality with delayed production of proinflammatory cytokines and chemokines required to resolve lung infections (8). Besides neutrophils, other leukocytes are important in resolving *A. baumannii* infections. Tissue-resident alveolar macrophages phagocytose *A. baumannii* before the recruitment of neutrophils to the lung, conferring an advantage during the early response to infection while neutrophils are being recruited (9). NK cells mediate pulmonary recruitment of neutrophils by producing the chemoattractant CXCL1, and depleting NK cells reduces neutrophil recruitment and impairs the clearance of *A. baumannii* from the lung (10). Besides the cell-mediated innate responses, studies have also reported the protective roles of the complement system and host-derived antimicrobial peptides (11, 12). Although *A. baumannii* clinical strains can completely resist complement-mediated lysis, opsonization with complement proteins facilitates the phagocytosis of multidrug-resistant *A. baumannii* by murine macrophages (13).

Whereas innate phagocyte responses against *A. baumannii* have been explored, the role of B and T cells is poorly understood. In a previous study, we reported a protective function of antibodies against *A. baumannii* in naive mice using a lymphocyte-deficient (*Rag2^-/-^*) mouse model. Presently, we demonstrate that B cells play a major protective role against *A. baumannii* as early as 4 hours post-infection (hpi). Notably, our finding indicates that B cells are necessary to prevent early extrapulmonary dissemination of *A. baumannii* and that mechanisms may not be strictly antibody-dependent. We further observed that B cell deficiency in mice results in pulmonary eosinophilia and increased accumulation of Ly6C^+^ NK cells in the lungs.

## Materials and Methods

### Mouse strains

C57BL/6NTac mice were obtained from Taconic Biosciences (Cambridge, Indiana). *Rag2^−/−^* (B6.Cg-Rag2^tm1.1Cgn^/J) and µMT (B6.129S2-Ighm^tm1Cgn^/J) mice were obtained from The Jackson Laboratory. All mice used in experiments were bred within our specific pathogen-free (SPF) facility, were 10 to 15 weeks of age, and sex-matched to their respective controls. All animal infections and procedures were performed in accordance with protocols approved by the Miami University Institutional Animal Care and Use Committee.

### Lung infection model

All mouse infections were performed using the multidrug-resistant *Acinetobacter baumannii* AB5075 clinical isolate (14). Bacteria were routinely cultured overnight (12 to 16 hours) in Luria-Bertani (LB) broth or on LB agar plates. Overnight cultures of *A. baumannii* strain AB5075 were diluted 1/1,000 in LB and grown at 37°C until reaching the mid-logarithmic phase (OD_600_ 0.4) with shaking at 200 rpm. Bacterial cells were collected by centrifugation, washed, and resuspended in sterile phosphate-buffered saline (PBS). Mice were anesthetized using 4% isoflurane and 2% oxygen, followed by intranasal inoculation with 1 × 10^8^ colony-forming units (CFU) suspended in 40 μL PBS. At indicated time points after infection, lungs, livers, and spleens were aseptically harvested and homogenized in 1 mL PBS using a minibead beater (BioSpec) for 25 seconds at 4,200 rpm. To determine bacterial loads, tissue lysates were serially diluted, plated on LB agar containing 25 μg/mL of ampicillin, and incubated overnight at 37°C.

### RNA sequencing and data analysis

RNA was extracted from whole lung tissue of uninfected WT (n=4), infected WT (n=4), uninfected µMT (n=3), and infected µMT (n=4) mice. Lung tissues were homogenized in 1 ml TRIzol reagent (Invitrogen) using a mini-bead beater, and total RNA was extracted using Direct-zol RNA Microprep Kit (Zymo Research) according to the manufacturer’s instructions. Library preparation was performed using the PerkinElmer NEXTFLEX Rapid Directional RNA-Seq Kit 2.0 with poly(A) captured mRNA. RNA sequencing was conducted using the NovaSeq 6000 System with S4 flow cells. RNA-Seq data were analyzed using the CLC Genomics Workbench 23.0.3 (Qiagen Bioinformatics) to identify differentially expressed genes. Low-quality reads were eliminated through trimming, and the high-quality reads were aligned to the *Mus musculus* sequence mm10 using default parameters, which allowed a maximum of 2 mismatches, setting a minimum length and similarity fraction of 0.8, and requiring a minimum of 10 hits per read. In this study, genes were considered differentially expressed based on the following criteria: false discovery rate (FDR) *P*-value ≤0.05 and absolute fold change ≥1.5. Raw datasets are available from the NCBI GEO database under accession: GSE267894.

### Flow cytometry of lung tissues

To prepare single-cell suspensions, lung tissue was excised and cut into small pieces in 7- mL Bijou containers (Greiner Bio-One) containing 1 mL Hanks’ balanced salt solution and 2.5 mg/mL of collagenase type 2 (Worthington Biochemical). Following incubation for 30 minutes at 37°C, lung homogenates were passed through 70-μm mesh filters. After centrifugation at 300 x *g* for 5 minutes, cells were resuspended in fluorescence-activated cell sorting buffer (PBS with 2% fetal bovine serum and 1 mM EDTA). Single-cell suspensions were stained for 30 minutes on ice with antibodies targeting the following surface proteins: CD45 (30-F11), CD11b (M1/70), F4/80 (BM8), Ly6C (HK1.4), Ly6G (1A8), Siglec-F (S170072), CD19 (6D5), CD3 (17A2) and NK 1.1 (PK136). Cell viability was measured using Zombie Aqua dye (Biolegend). Data acquisition was performed using an Attune NxT flow cytometer (ThermoFisher). Results were analyzed using FlowJo software version 10.8.2.

### Isolation of mouse NK cells

Single-cell suspension from mouse lung tissues was prepared as described above. NK cells were first enriched by negative selection using MojoSort Mouse NK Cell Isolation Kit (BioLegend) according to the manufacturer’s instructions. To increase the purity, NK cells from negative selection were further enriched by positive selection using biotinylated anti-mouse NK1.1 antibody (Miltenyi Biotec). Isolated NK cells were directly lysed in Trizol reagent and total RNA was extracted for RT-qPCR.

### RT-qPCR

cDNA was prepared from RNA by reverse transcription (RT) using the iSCRIPT cDNA synthesis kit (Bio-Rad). Quantitative polymerase chain reaction (qPCR) was performed using a CFX connect real-time PCR detection system (Bio-Rad) with the following Thermo Fisher Scientific Taqman probes: Mm04207460_m1 (*Cxcl1*), Mm00442837_m1 (*Gzmb*), Mm03807119_mH (*Klra8*), Mm01168134_m1 (*Ifng*) and Mm04394036_g1 (*Actb*).

### Statistical Analyses

Statistical analyses for non-sequencing experiments were performed using GraphPad Prism version 10.2.0. *P* values ≤ 0.05 were considered statistically significant in these analyses.

## Results

### Innate B cells are crucial for protection against *A. baumannii* infection

In a previous study, we reported that *Rag2^-/-^* mice lacking both B and T cells were more susceptible to *A. baumannii* infection than WT mice, and prophylactic administration of serum or normal immunoglobulins could compensate for B and T cell deficiency (13). However, it remained possible that the lack of T cells could contribute to increased susceptibility of *Rag2^-/-^* mice at 24 hpi, since γδ and NK-T cells are protective against bacterial pneumonia (15, 16). Do determine the specific contribution of B cells to *A. baumannii* resistance, we compared the bacterial load among WT, *Rag2^-/-^*, and B cell-deficient mice (µMT) mice at 24 hpi. The µMT mice had a significantly higher bacterial load in lung, liver, and spleen than WT mice. Notably, the bacterial load between µMT and *Rag2^-/-^* mice was comparable (Figure 1). These observations indicate that innate B cell functions are crucial in protecting against *A. baumannii* respiratory infection at 24 hpi.

**Figure 1.**
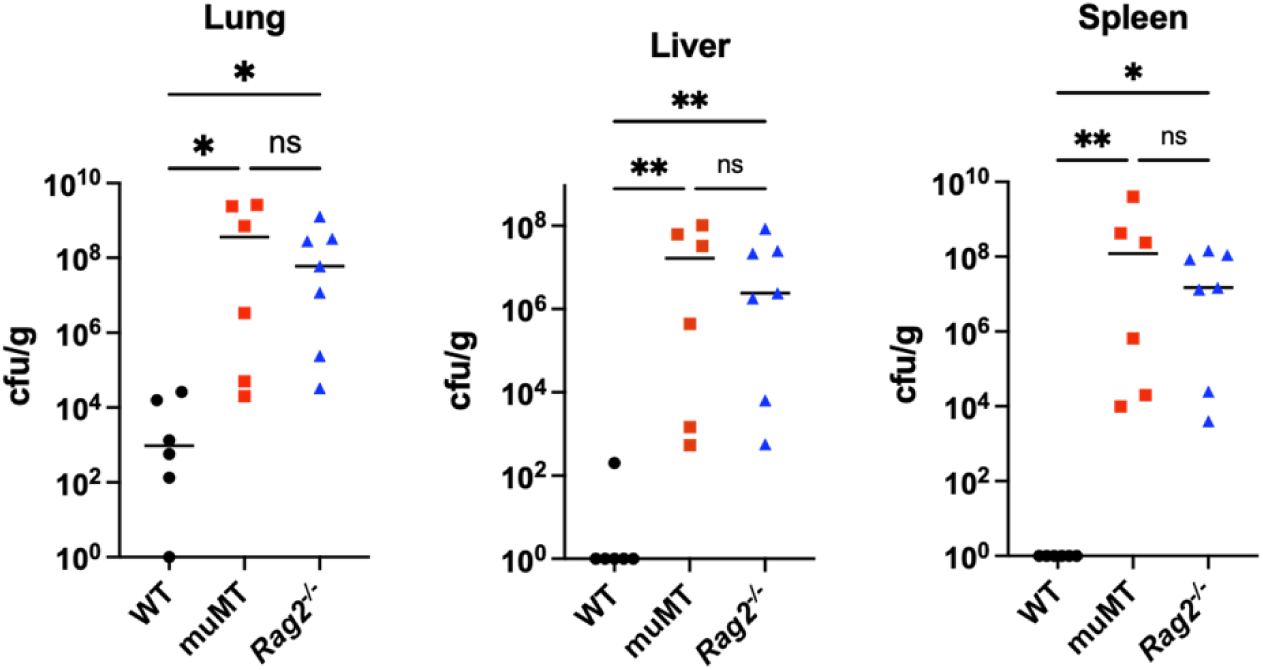
Bacterial load at 24 hpi. Mice were intranasally infected with 10^8^ CFU of *A. baumannii* strain AB5075. Bacterial burden in lung, liver, and spleen was measured at 24 hpi. Data shown are combined from 2 independent experiments. Each symbol represents data from a single mouse. Statistical analyses were performed by the Kruskal-Wallis test (*, *P* ≤ 0.05; **, *P* ≤ 0.01) and horizontal bars represent the median of each data set.

### B cell deficiency results in increased systemic dissemination of *A. baumannii* early after infection

While B cell-deficient mice are highly bacteremic by 24 hours after infection, early responses are likely to be key to bacterial resistance. *A. baumannii* can induce a very strong pulmonary inflammatory response, and a previous study suggests that neutrophils are recruited to the lung as early as 4 hours post-infection (8). Therefore, we also measured the bacterial load at 4 hpi to determine if there was any significant difference in bacterial load between WT and µMT mice at an early time point. We did not observe any significant difference in lung bacterial load between WT and µMT mice at 4 hpi (Figure 2). However, the livers and spleens of µMT mice had significantly higher bacterial burden compared to WT mice (Figure 2). These findings indicate increased systemic dissemination of *A. baumannii* in µMT mice as early as 4 hpi. This suggests that the severe bacteremia observed at 24 hpi may result from either early escape from the lung or failure to control disseminated organisms.

**Figure 2.**
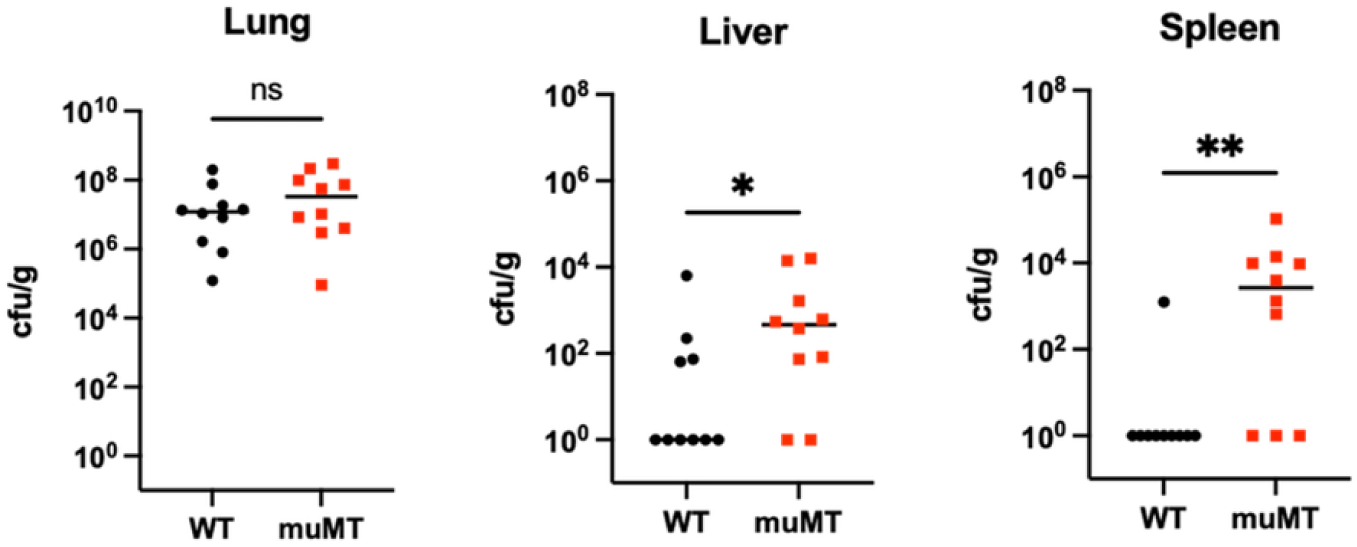
Bacterial load at 4 hpi. Mice were intranasally infected with 10^8^ CFU of *A. baumannii* strain AB5075. Bacterial burden in lung, liver, and spleen was measured at 4 hpi. Data shown are combined from 3 independent experiments. Each symbol represents data from a single mouse. Statistical analyses were performed by the Mann-Whitney U test (*, *P* ≤ 0.05; **, *P* ≤ 0.01; ns, not significant), and horizontal bars represent the median of each data set.

### Transcriptomic analysis of *A. baumannii*-infected lungs

The increased extrapulmonary dissemination of *A. baumannii* in µMT mice at 4 hpi indicates that the µMT phenotype could be associated with a qualitatively different inflammatory response compared to WT mice. Therefore, to better understand immune responses against *A. baumannii* in lungs, we performed RNA-sequencing (RNA-Seq) on whole lung tissue isolated from WT and µMT mice. We first compared the gene expression profiles between infected (at 4 hpi) and uninfected WT mice. Overall, there were 4,120 differentially expressed genes (DEGs), with 1,987 genes upregulated and 2,133 down-regulated by infection (fold change, ≥1.5; FDR *P*- value <.05). A heatmap of the 50 most differentially expressed genes in response to infection is shown in Figure 3. It includes many genes involved in proinflammatory cytokine responses. Top genes upregulated by *A. baumannii* infection include *Gm4070,* encoding an interferon-induced large-GTPase; *Cxcl2* and *Ccl20*, encoding important pro-inflammatory chemokines; *Csf3,* encoding G-CSF for phagocyte functional enhancement; and *Acod1,* encoding Aconitate decarboxylase 1, an important metabolic regulator of immune function. This shows that *A. baumannii* induces a strong inflammatory response in the lung within four hours of infection (Figure 3).

**Figure 3.**
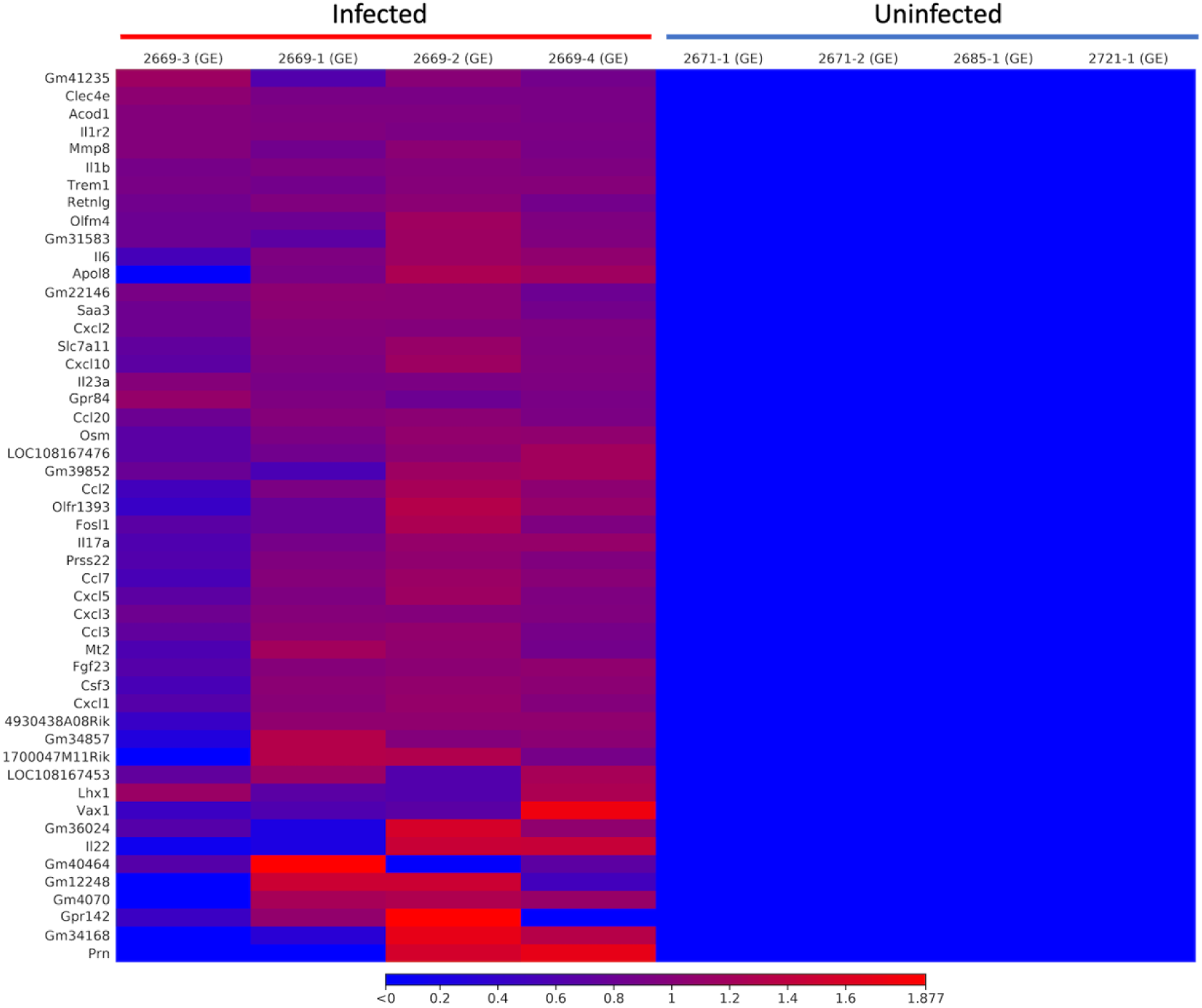
Heatmap showing the top 50 highly differentially expressed genes induced by *A. baumannii* at 4 hpi. Heatmap was generated using CLC genomics workbench software version 23.0.3. The color scale represents the expression level of the genes (log CPM).

We next compared the differential gene expression profiles of WT and µMT mice. As expected from a comparison where one mouse lacks B cells, the top differentially expressed genes between WT and µMT mice were directly related to B cell functions. Therefore, we manually filtered out genes with substantial expression by B cells (17). In naïve mice, this approach resulted in 182 genes that were more highly expressed in uninfected WT mice than uninfected µMT mice (fold change, ≥1.5; FDR *P*-value < 0.05). In addition to the absence of antibodies, Gene Ontology (GO) enrichment analysis of these genes suggests that naive WT mice likely have an increased antimicrobial capacity at homeostasis compared to µMT mice. Notably, genes encoding antimicrobial proteins such as *Bpifa1, Reg3g, Pglyrp1,* and *Ltf* all had constitutively higher levels of expression in the lungs of WT mice compared with µMT mice, potentially providing an antibody-independent mechanism of bacterial control that is impaired in the absence of B cells. In addition, genes involved in the chemotaxis of NK cells and neutrophils were also affected (Table 1).

**Table 1.**
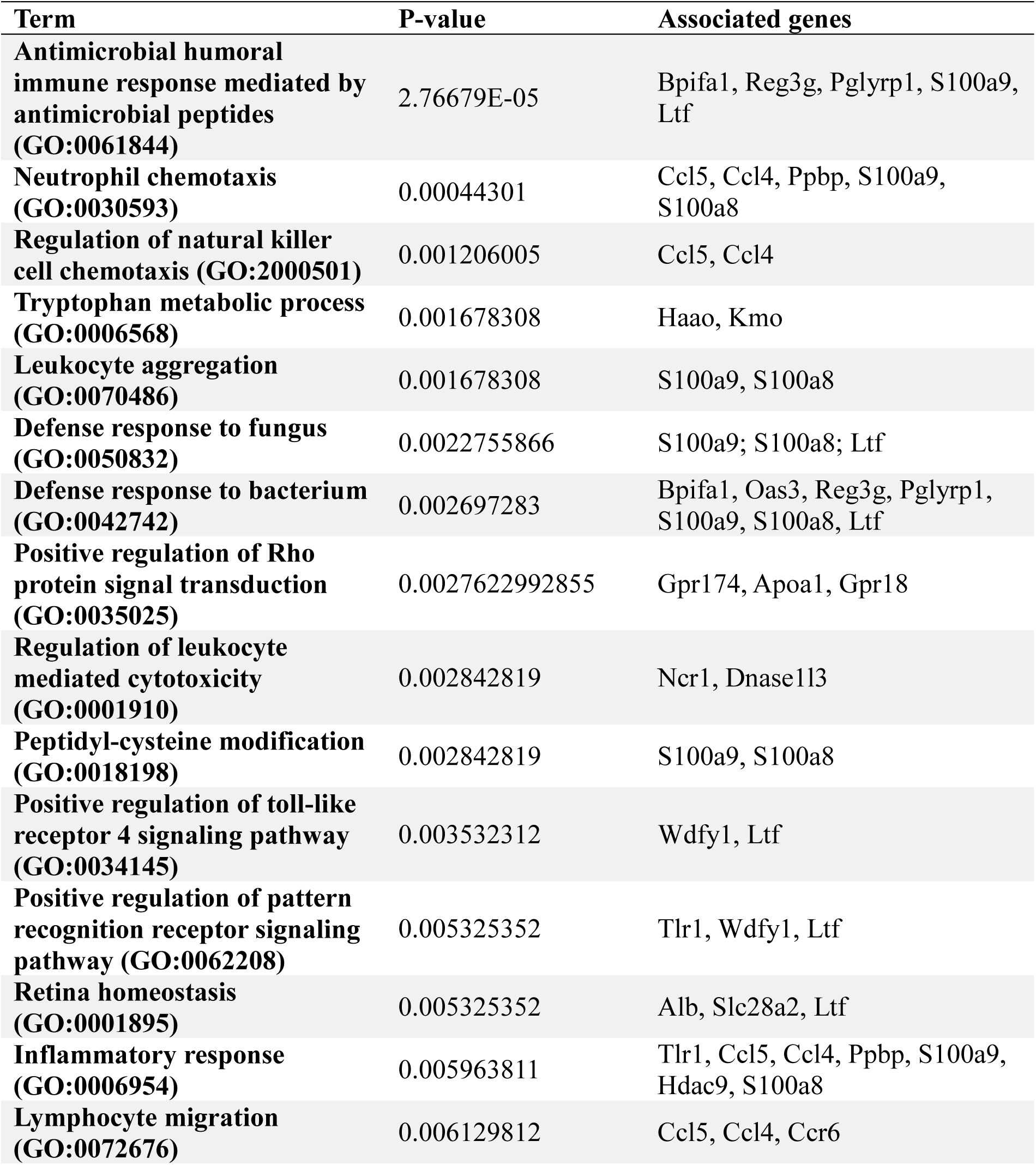
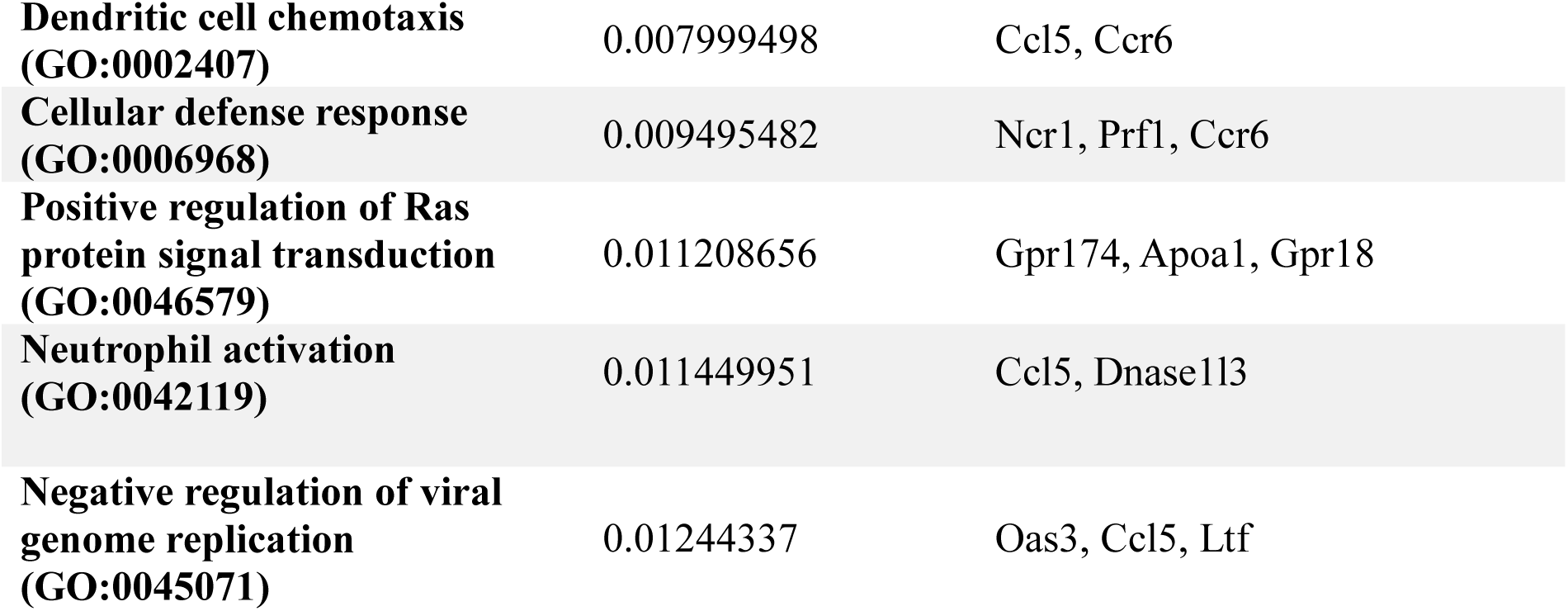
GO enrichment analysis using genes that were significantly highly expressed in naive WT mice compared to muMT mice. The top 20 enriched terms are shown in the table based on *P*-values (Redundant GO terms with the same gene set are excluded in this table). Analysis was done using Enrichr (https://amp.pharm.mssm.edu/Enrichr/).

Gene expression data were also examined after four hours of infection. To remove confounding signals, genes differentially expressed before infection were filtered from the list of differentially expressed genes. With these genes eliminated, 32 genes were more highly expressed in WT compared to µMT mice after infection (Supplementary Table 1). Despite early deviation in the response to bacterial infection between WT and µMT mice, GO enrichment analysis of this gene set failed to produce a coherent set of biological processes that might explain the resistance phenotype, perhaps indicating that the most important differences were present before infection. Similarly, after filtering out genes with differential expression before infection, 75 genes were more highly expressed by µMT lungs than by WT lungs. Using the Knock Out Mouse Phenotyping Program (KOMP2) libraries for these genes, GO analysis indicated abnormally high expression of genes with an annotated association with uterine hydrometra (*Bpifb1, Clca1, Prss35, Ccl28,* and *Gpld1*), perhaps indicating early problems with fluid accumulation in B cell-deficient lungs. Also with notably higher expression in µMT mice was *Asb2*, which is associated with the recruitment of Ly6C-positive NK cells (Supplementary Tables 2 and 3).

### B cell deficiency results in pulmonary eosinophilia

While not specified by Gene Ontology analysis, our transcriptomic data showed relatively higher expression of genes coding for CCL8 and CCL28, chemokines known to recruit eosinophils (18, 19), in the lungs of infected µMT mice (Supplementary Table 2). Hypothesizing that a gene expression signature indicating excess fluid accumulation, combined with increased extrapulmonary dissemination of *A. baumannii*, may be the result of cell-mediated tissue damage, flow cytometry analysis was performed to examine B cell-dependent recruitment of eosinophils and other inflammatory cell subsets. We first compared the frequency of neutrophils and eosinophils in the lungs in WT and µMT mice, both before and after infection. We did not see any significant difference in the frequency of neutrophils in the lungs of WT and µMT mice. We also specifically examined the frequency of Siglec-F expressing neutrophils in mouse lungs since Siglec-F^+^ neutrophils have pro-inflammatory activity and produce more neutrophil extracellular traps during bacterial infection (20). However, the frequency of Siglec-F^+^ neutrophils was comparable between the two groups of mice (Figure 4). In contrast, B cell deficiency resulted in a significantly increased frequency of pulmonary eosinophils in uninfected and infected conditions (Figure 4). This implicates B cells in eosinophil homeostasis in the lung.

**Figure 4.**
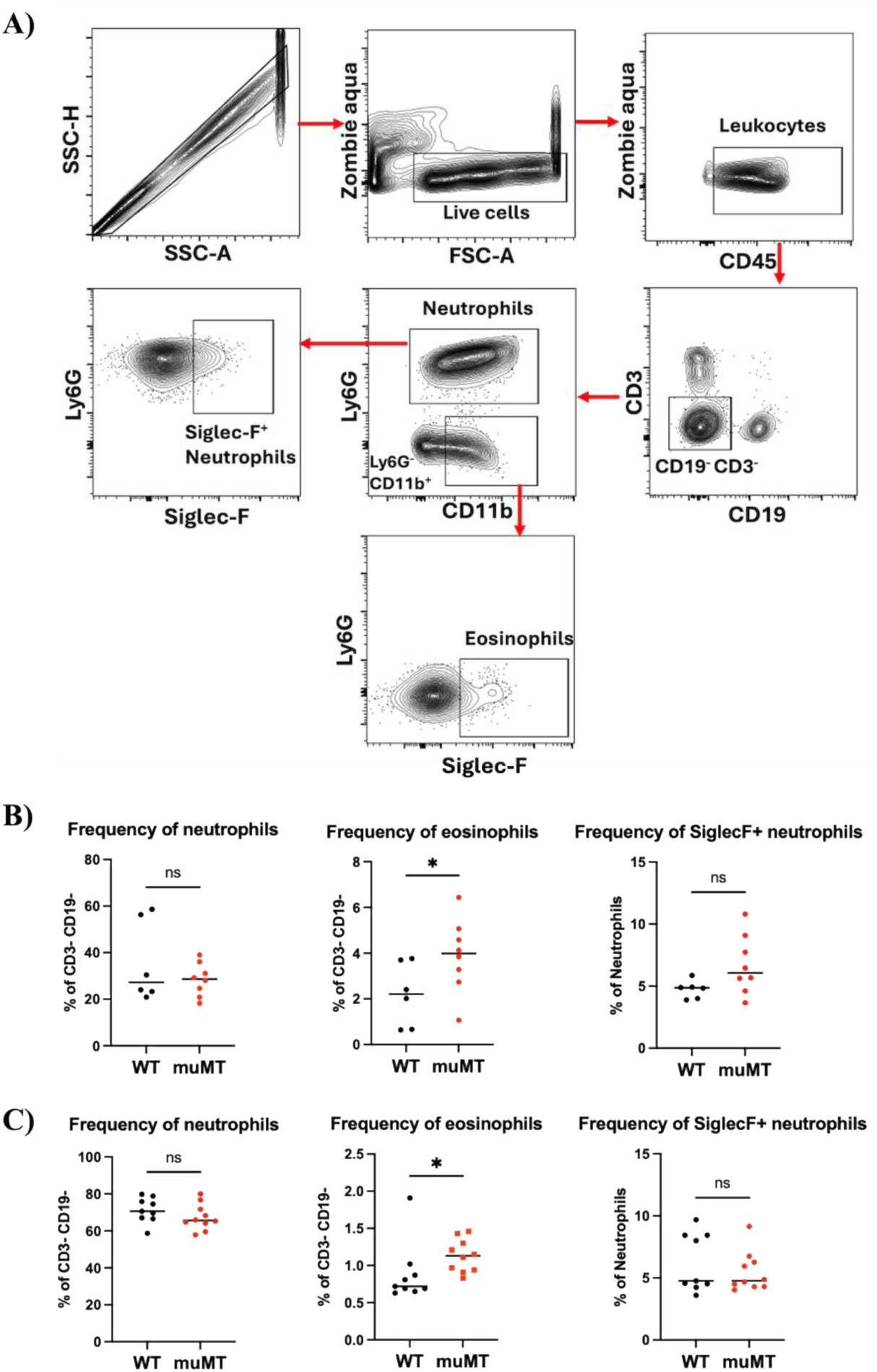
A) Gating strategy to analyze recruitment of immune cells by flow cytometry. **B)** Frequency of neutrophils, eosinophils, and Siglec-F^+^ neutrophils in lungs of uninfected WT and muMT mice. **C)** Frequency of neutrophils, eosinophils, and SiglecF^+^ neutrophils in lungs of WT and muMT mice at 4 hpi. Data shown are combined from at least 2 independent experiments. Statistical analyses were performed by the Mann-Whitney U test (*, *P* ≤ 0.05; ns, not significant) and horizontal bars represent the median of each data set.

### B cell-deficient mice had increased frequency of pulmonary Ly6C+ NK cells at 4 hpi

Our lung transcriptomic data indicate that B cell deficiency may regulate the recruitment of Ly6C^+^ NK cells (Table 3). A previous study also reported an increased recruitment of NK cells to the lung tumor microenvironment in µMT mice compared to WT mice, which was attributed to an elevated number of type I interferon-producing plasmacytoid dendritic cells in µMT mice (21). To determine any difference in the recruitment of pulmonary NK cells between WT and µMT mice, we analyzed lung tissues by flow cytometry. While we did not observe any significant difference in the frequency of NK cells in the lungs of WT and µMT mice (Figure 5B and 5C), we observed that NK cells in WT and µMT mice differed in their expression of Ly6C at 4 hpi. WT mice had a reduced frequency of Ly6C^+^ NK cells compared to µMT mice (Figure 5C). Notably, this difference in the frequency of Ly6C^+^ NK cells was not observed in uninfected mice, indicating that *A. baumannii* infection facilitates the differential recruitment and/or Ly6C expression among NK cells in WT and µMT mice. A prior study suggests that Ly6C^neg^ NK cells are involved in increased production of interferon-gamma and granzyme B compared to Ly6C^+^ NK cells (22). To determine if there are any functional differences in NK cells between WT and µMT mice, we isolated mouse NK cells from lungs by magnetic enrichment at 4 hpi. We measured by RT-qPCR the expression of four genes (*Ifng, Gzmb, Klra8, and Cxcl1)* expressed in activated NK cells. However, we did not observe any significant difference in the expression of these genes in NK cells between WT and µMT mice (Supplementary Figure 1), implying that differences in Ly6C expression may indicate functional differences apart from gamma interferon production in the lung NK cell populations.

**Figure 5.**
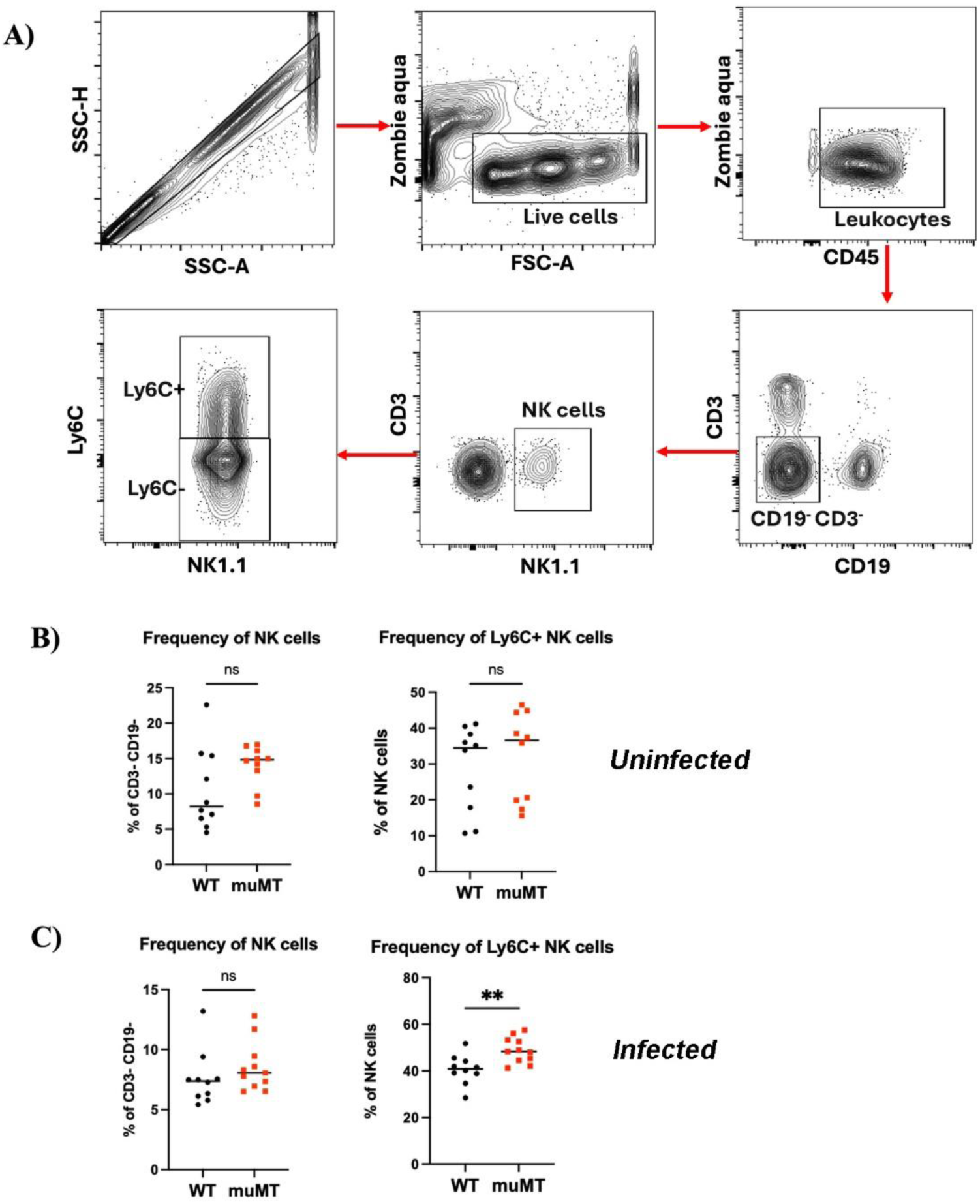
A) Gating strategy to analyze lung NK cells. **B**) Frequency of NK cells and Ly6C^+^ NK cells in lungs of uninfected mice. **C**) Frequency of NK cells and Ly6C^+^ NK cells at 4 hpi. Data shown are combined from 3 independent experiments. Each symbol represents data from a single mouse. Statistical analyses were performed by the Mann-Whitney U test (**, *P* ≤ 0.01; ns, not significant) and horizontal bars represent the median of each data set.

## Discussion

Our previous study found that natural antibodies could provide innate resistance to A. baumannii lung infection but left open the possibility that the susceptibility of Rag2-/- mice to A. baumannii could be explained by a deficiency in innate T cells (13). Therefore, in the present study, we compared the susceptibility of WT, Rag2-/-, and B cell-deficient mice (µMT) to A. baumannii. Since the bacterial load between µMT and Rag2-/- mice were comparable, our data indicated that innate B cells are crucial in protecting against A. baumannii infection at 24 hpi. While the lung bacterial load in WT and µMT mice was comparable at 4 hpi, µMT mice exhibited significantly higher extrapulmonary dissemination, demonstrating that B cells are critical for providing early protection from breaching of the epithelium and resulting septicemia.

Whether the difference between WT and µMT mice is the principal result of reduced antimicrobial proteins, impaired lung macrophage and NK cell effector functions, and/or antibody-dependent responses in circulation remains to be determined. Our RNA-seq data suggest that highly expressed genes (unrelated to canonical B cell function) in naive WT mice involve antimicrobial peptide production, neutrophil chemotaxis, neutrophil activation, and NK cell-mediated responses. More specifically, several genes that code for proteins with antimicrobial activity had significantly higher expression in naive WT mice compared to µMT mice (23–26). BPIFA1 (BPI fold-containing group A member 1) is an airway host-protective protein that binds to bacterial LPS and has direct bactericidal activity (23). This protein can also regulate lung neutrophil recruitment during acute inflammation (27). The calprotectin complex encoded by *S100a8* and *S100a9* inhibits bacterial growth by sequestering manganese (Mn) and zinc (Zn), which are essential metals for the growth of *A. baumannii* in murine pneumonia (28). Additionally, calprotectin is involved in neutrophil migration to inflammatory sites (26, 29). PGLYRP1 (peptidoglycan recognition protein 1) interferes with peptidoglycan biosynthesis and is directly bactericidal (30). Therefore, higher expression of genes encoding the antimicrobial peptides may facilitate increased clearance of bacteria in WT mice. Some antimicrobial peptides, including calprotectin, PGLYRP1, and LTF (lactoferrin), are produced by activated neutrophils (30–32), leading us to speculate that antibodies enhance their production by infiltrating neutrophils. Overall, the higher expression of genes related to antimicrobial peptide production and immune cell recruitment in naive WT mice suggests that B cells or their antibody products are necessary for innate antimicrobial responses.

In addition to reduced innate antimicrobial functions, B cell deficiency increased the frequency of pulmonary eosinophils. This is consistent with a separate study that reported B cell deficiency results in allergic eosinophil-rich inflammation in the lungs and airways (33). This increased number of eosinophils may contribute to enhanced pulmonary inflammation and tissue damage, ultimately leading to systemic dissemination of bacteria (34). Although it is not clear why B cell-deficient mice exhibited an increased frequency of eosinophils, one possible explanation could be the lack of B regulatory cells (Bregs). Bregs have immunosuppressive capacity, often mediated by IL-10 secretion, and can suppress eosinophil-mediated airway allergic responses (35, 36). This raises the possibility that B1a cells, which are a source of natural antibodies and also produce a substantial amount of IL-10, play a crucial role against *A. baumannii* infection, as both natural antibodies and IL-10 are protective against *A. baumannii* infection (13, 37, 38). Therefore, further investigation of the role of innate B cells in controlling allergic lung inflammation during bacterial infection is warranted.

Our study also found that genes (*Ncr1, Prf, Gzmb, Nkg7,* and *Klra8*) associated with NK cell-mediated functions were more highly expressed in naive WT mice than in µMT mice. It is unclear that NK cells could be directly involved in clearing *A. baumannii*; however, NK cells may play a role in bacterial clearance through interactions with resident or recruited phagocytes. We speculated that differences in NK cell activation might alter macrophage activation and function through IFNγ. WT mice have a higher frequency of pulmonary Ly6C^-^ NK cells compared to µMT mice at 4 hpi, and a previous study reported that Ly6C^-^ NK cells are involved in increased production of interferon-gamma and granzyme B compared to Ly6C^+^ NK cells (22). However, we did not observe any significant difference in the expression of activation-associated genes in NK cells between WT and µMT mice. Although our data indicate that the total NK cells between WT and µMT mice might have a similar level of expression of the genes tested in this study, we cannot rule out the possibility that Ly6C^+^ and Ly6C^-^ subsets of NK cells may differ functionally (39–41). Therefore, the novel findings from our study related to the role of B cells in recruiting different subsets of NK cell population warrants further investigation to understand how different subsets of NK cell population are regulated by B cells during *A. baumannii* pulmonary infection.

In summary, this study suggests that B cells or their antibody products regulate the recruitment and activation of other innate immune cells, including eosinophils and NK cells, preventing systemic dissemination of *A. baumannii*. Our study is the first to describe significantly reduced expression of genes that code for antimicrobial peptides in B cell-deficient mice. This finding highlights the importance of studying innate B cells and natural antibodies in maintaining an antimicrobial response in the absence of infection. Such heightened antimicrobial responses could significantly impact innate immune resistance when the host is exposed to pathogens. The novel findings related to B cell-mediated function increase our understanding of the role of innate lymphocytes in infection. Additional investigation will provide a more complete understanding of how B cells regulate antimicrobial activity, even in the absence of acute infection, which may aid future research in developing effective immunotherapies.

## Acknowledgements

The Rush University Genomics and Microbiome Core Facility provided RNA sequencing support. The authors thank Dr. Andor Kiss and the Center for Bioinformatics and Structural Genomics at Miami University for instrumentation and computational support. Support was provided in-part by grants from the National Institute for Allergy and Infectious Diseases at the National Institutes of Health [grant numbers R15AI138184 and R15AI174170 to TW] and the Miami University Committee for Faculty Research.

## Author contributions

AI performed experiments and wrote and edited the manuscript; LAA provided critical reagents, directed research, and edited the manuscript; TJW directed research and edited the manuscript.

---

**Supplementary Figure 1.**
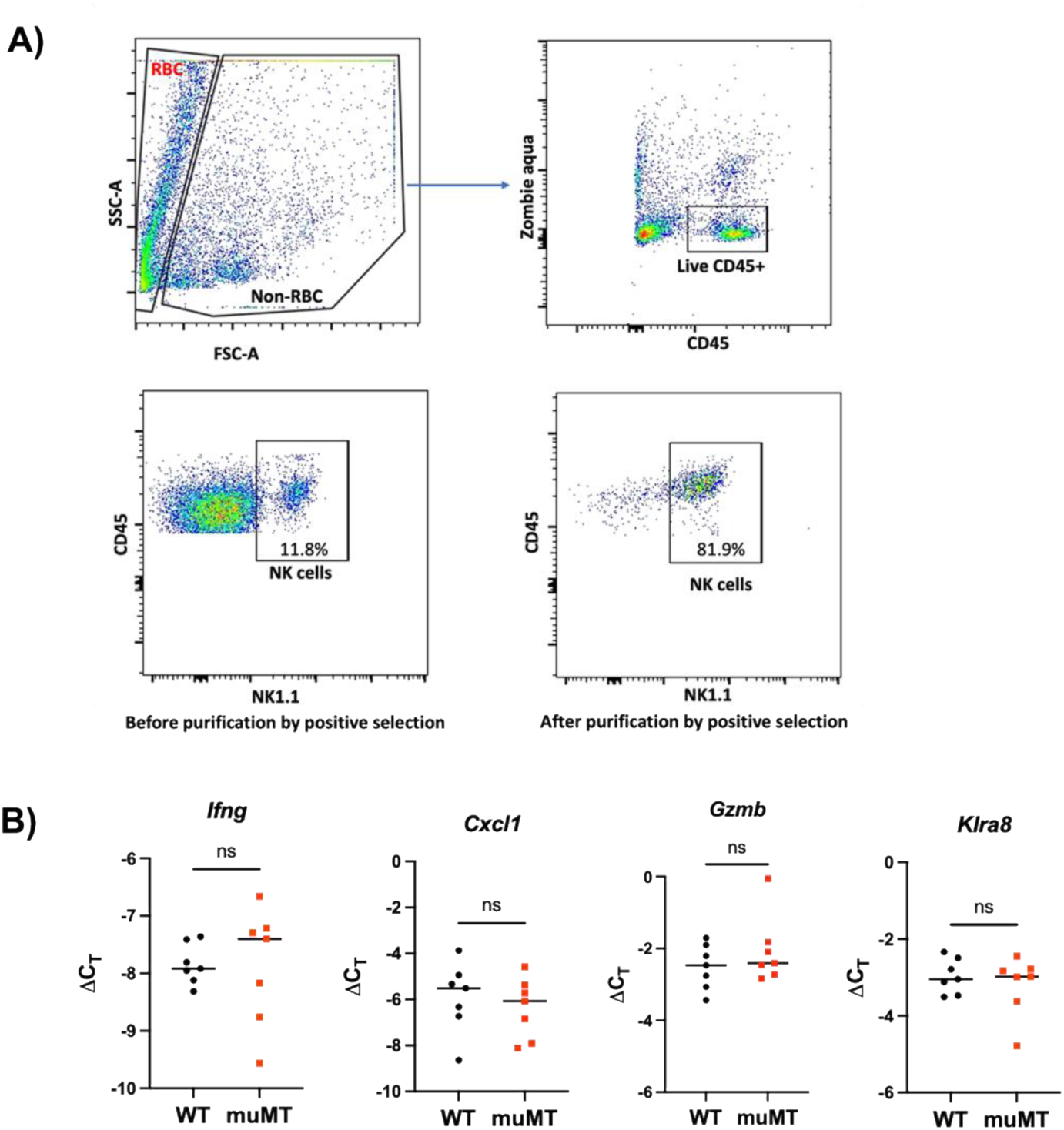
A) Gating strategy to check the purity of isolated NK cells. NK cells from lung tissue were first purified through negative selection using the MojoSort kit from BioLegend. NK cells were further enriched by positive selection using biotinylated anti-mouse NK1.1 antibody. A representative flow plot is shown. The purity of the isolated NK cells ranged from 60% to 86%. B) Expression of *Ifng*, *Cxcl1*, *Gzmb*, and *Klra8* in NK cells tested by RT-qPCR. Delta Ct values were calculated by subtracting the Ct value of the target gene from the Ct value of the beta-actin gene, which was used as a constitutively expressed gene control. Data shown are combined from 2 independent experiments. Each symbol represents data from a single mouse. Statistical analyses were performed by the Mann-Whitney U test (ns, not significant), and horizontal bars represent the median of each data set.

**Supplementary Table 1.** – Genes with higher expression in WT mice compared with μMT mice 4 hours after *A. baumannii* infection (FDR p<0.05, fold-change ≥ 1.5).

| Gene | Max group mean | Log<sub>2</sub> fold change | Fold change | FDR p-value | GeneID |
| --- | --- | --- | --- | --- | --- |
| **Pira2** | 22.1 | 1.39 | 2.6 | $5.096 \times 10^{-11}$ | 18725 |
| **Plat** | 108.5 | 0.81 | 1.8 | $5.196 \times 10^{-7}$ | 18791 |
| **Dusp8** | 34.1 | 0.88 | 1.8 | $1.224 \times 10^{-4}$ | 18218 |
| **Gdnf** | 8.5 | 0.88 | 1.8 | $1.597 \times 10^{-4}$ | 14573 |
| **Bhlhe41** | 17.1 | 0.63 | 1.5 | $2.049 \times 10^{-4}$ | 79362 |
| **Ccl20** | 583.5 | 0.99 | 2.0 | $2.491 \times 10^{-4}$ | 20297 |
| **Pbxip1** | 102.4 | 0.95 | 1.9 | $4.407 \times 10^{-4}$ | 229534 |
| **LOC102634296** | 0.6 | 3.04 | 8.2 | $5.224 \times 10^{-4}$ | 102634296 |
| **Dbp** | 143.6 | 0.76 | 1.7 | $6.049 \times 10^{-4}$ | 13170 |
| **Snca** | 36.8 | 0.83 | 1.8 | $6.972 \times 10^{-4}$ | 20617 |
| **Zcchc18** | 1.3 | 1.54 | 2.9 | $1.360 \times 10^{-3}$ | 66995 |
| **Gm40525** | 2.7 | 2.85 | 7.2 | $1.678 \times 10^{-3}$ | 105245011 |
| **Cdr2** | 51.3 | 0.61 | 1.5 | $2.808 \times 10^{-3}$ | 12585 |
| **Hbb-bt** | 3035.0 | 0.73 | 1.7 | $3.861 \times 10^{-3}$ | 101488143 |
| **Pla1a** | 6.9 | 0.83 | 1.8 | $4.112 \times 10^{-3}$ | 85031 |
| **Ptpn22** | 7.0 | 0.72 | 1.6 | $4.579 \times 10^{-3}$ | 19260 |
| **Vipr2** | 3.7 | 0.73 | 1.7 | $4.579 \times 10^{-3}$ | 22355 |
| **Klf4** | 337.3 | 0.60 | 1.5 | $5.319 \times 10^{-3}$ | 16600 |
| **Fam71f2** | 5.0 | 0.70 | 1.6 | $1.131 \times 10^{-2}$ | 245884 |
| **8030474K03Rik** | 0.4 | 3.30 | 9.9 | $1.352 \times 10^{-2}$ | 382231 |
| **Tnfrsf10b** | 11.4 | 0.72 | 1.7 | $1.352 \times 10^{-2}$ | 21933 |
| **Gm40117** | 3.7 | 0.93 | 1.9 | $1.423 \times 10^{-2}$ | 105244523 |
| **Nr4a1** | 59.3 | 0.78 | 1.7 | $1.581 \times 10^{-2}$ | 15370 |
| **Nr1d1** | 101.7 | 0.73 | 1.7 | $1.806 \times 10^{-2}$ | 217166 |
| **Cftr** | 5.4 | 0.73 | 1.7 | $1.878 \times 10^{-2}$ | 12638 |
| **A430078G23Rik** | 2.6 | 1.00 | 2.0 | $1.906 \times 10^{-2}$ | 319493 |
| **Gm19696** | 0.9 | 1.16 | 2.2 | $2.233 \times 10^{-2}$ | 100503447 |
| **Acox1** | 16.8 | 0.59 | 1.5 | $2.541 \times 10^{-2}$ | 74121 |
| **Gm17619** | 1.3 | 1.47 | 2.8 | $2.651 \times 10^{-2}$ | 100502923 |
| **LOC105244034** | 4.1 | 0.97 | 2.0 | $2.948 \times 10^{-2}$ | 105244034 |
| **Gm42078** | 0.8 | 3.71 | 13.0 | $3.268 \times 10^{-2}$ | 105246872 |
| **Rspo4** | 1.3 | 1.14 | 2.2 | $3.268 \times 10^{-2}$ | 228770 |

**Supplementary Table 2:** Genes with higher expression in lungs of μMT mice compared to WT mice 4 hours after *A. baumannii* infection.

| Gene | Max<br>group<br>mean | Log<sub>2</sub><br>fold<br>change | Fold<br>change | FDR p-value | GeneID |
| --- | --- | --- | --- | --- | --- |
| G530011O06Rik_1 | 5.6 | 2.51 | 5.7 | 6.2*10<sup>-8</sup> | 654820 |
| Bpifb1 | 144.5 | 0.69 | 1.6 | 2.3*10<sup>-6</sup> | 228801 |
| Gm7120 | 5.1 | 1.36 | 2.6 | 2.6*10<sup>-6</sup> | 633640 |
| Ccr2 | 34.9 | 1.33 | 2.5 | 2.8*10<sup>-6</sup> | 12772 |
| Ttn | 0.9 | 0.96 | 1.9 | 2.8*10<sup>-5</sup> | 22138 |
| Eef1a2 | 15.0 | 0.99 | 2.0 | 3.1*10<sup>-5</sup> | 13628 |
| Myoz2 | 24.7 | 0.96 | 1.9 | 5.8*10<sup>-5</sup> | 59006 |
| Sept3 | 3.5 | 0.94 | 1.9 | 7.3*10<sup>-5</sup> | 24050 |
| Clca1 | 2.7 | 1.79 | 3.5 | 2.2*10<sup>-4</sup> | 23844 |
| Cyp1a1 | 134.4 | 2.88 | 7.4 | 2.3*10<sup>-4</sup> | 13076 |
| Gkn3 | 6.5 | 1.56 | 2.9 | 2.3*10<sup>-4</sup> | 68888 |
| Cmya5 | 2.5 | 0.81 | 1.8 | 2.5*10<sup>-4</sup> | 76469 |
| Nppa | 63.3 | 1.36 | 2.6 | 2.9*10<sup>-4</sup> | 230899 |
| F13a1 | 38.1 | 1.35 | 2.6 | 3.1*10<sup>-4</sup> | 74145 |
| Fmod | 15.1 | 0.83 | 1.8 | 5.0*10<sup>-4</sup> | 14264 |
| Ckm | 25.0 | 0.79 | 1.7 | 5.4*10<sup>-4</sup> | 12715 |
| Tmc5 | 3.0 | 1.07 | 2.1 | 6.2*10<sup>-4</sup> | 74424 |
| Cd207 | 3.3 | 1.32 | 2.5 | 6.3*10<sup>-4</sup> | 246278 |
| Prss35 | 1.1 | 1.45 | 2.7 | 7.9*10<sup>-4</sup> | 244954 |
| Mb | 90.6 | 0.82 | 1.8 | 8.5*10<sup>-4</sup> | 17189 |
| Actc1 | 278.1 | 0.72 | 1.6 | 1.4*10<sup>-3</sup> | 11464 |
| Mafb | 15.5 | 1.04 | 2.1 | 1.5*10<sup>-3</sup> | 16658 |
| Synb | 0.4 | 2.23 | 4.7 | 2.3*10<sup>-3</sup> | 239167 |
| Cygb | 63.0 | 0.60 | 1.5 | 2.6*10<sup>-3</sup> | 114886 |
| Rps27 | 80.6 | 2.42 | 5.4 | 3.0*10<sup>-3</sup> | 57294 |
| Tifab | 6.3 | 0.96 | 1.9 | 3.0*10<sup>-3</sup> | 212937 |
| Myl3 | 11.9 | 1.20 | 2.3 | 3.2*10<sup>-3</sup> | 17897 |
| Kcnj3 | 1.1 | 1.26 | 2.4 | 3.4*10<sup>-3</sup> | 16519 |
| Ms4a6c | 11.8 | 0.94 | 1.9 | 3.8*10<sup>-3</sup> | 73656 |
| Ldb3 | 6.0 | 0.76 | 1.7 | 4.0*10<sup>-3</sup> | 24131 |
| Asb2 | 5.8 | 0.86 | 1.8 | 4.5*10<sup>-3</sup> | 65256 |
| Ms4a4a | 12.2 | 1.30 | 2.5 | 4.6*10<sup>-3</sup> | 666907 |
| Lmod2 | 9.4 | 0.84 | 1.8 | 4.9*10<sup>-3</sup> | 93677 |
| Myom2 | 5.3 | 0.90 | 1.9 | 5.1*10<sup>-3</sup> | 17930 |
| Sfrp4 | 1.9 | 1.33 | 2.5 | 6.1*10<sup>-3</sup> | 20379 |
| Il11ra2 | 1.1 | 1.65 | 3.1 | 9.0*10<sup>-3</sup> | 16158 |
| Myh6 | 145.4 | 0.73 | 1.7 | 1.0*10<sup>-2</sup> | 17888 |
| **Myl4** | 67.6 | 0.64 | 1.6 | $1.0 \cdot 10^{-2}$ | 17896 |
| **Obscn** | 1.3 | 0.73 | 1.7 | $1.1 \cdot 10^{-2}$ | 380698 |
| **Gm21104** | 14.9 | 1.19 | 2.3 | $1.1 \cdot 10^{-2}$ | 100861647 |
| **Scara5** | 5.4 | 0.63 | 1.5 | $1.1 \cdot 10^{-2}$ | 71145 |
| **Gcsam** | 1.6 | 1.06 | 2.1 | $1.1 \cdot 10^{-2}$ | 14525 |
| **Gpld1** | 4.5 | 0.80 | 1.7 | $1.2 \cdot 10^{-2}$ | 14756 |
| **Hmcn2** | 0.9 | 0.81 | 1.8 | $1.2 \cdot 10^{-2}$ | 665700 |
| **Fcgr1** | 4.9 | 1.18 | 2.3 | $1.2 \cdot 10^{-2}$ | 14129 |
| **LOC108168091** | 0.5 | 3.46 | 11.0 | $1.3 \cdot 10^{-2}$ | 108168091 |
| **Gpc6** | 1.0 | 0.74 | 1.7 | $1.4 \cdot 10^{-2}$ | 23888 |
| **Itgb8** | 0.4 | 2.27 | 4.8 | $1.4 \cdot 10^{-2}$ | 320910 |
| **Nccrp1** | 3.0 | 1.21 | 2.3 | $1.4 \cdot 10^{-2}$ | 233038 |
| **Cldn1** | 8.7 | 0.69 | 1.6 | $1.4 \cdot 10^{-2}$ | 12737 |
| **Ccl8** | 5.6 | 1.38 | 2.6 | $1.4 \cdot 10^{-2}$ | 20307 |
| **Muc5b** | 15.2 | 0.76 | 1.7 | $1.5 \cdot 10^{-2}$ | 74180 |
| **Casq2** | 9.0 | 0.95 | 1.9 | $1.5 \cdot 10^{-2}$ | 12373 |
| **Hrc** | 7.3 | 0.85 | 1.8 | $1.6 \cdot 10^{-2}$ | 15464 |
| **Msr1** | 3.6 | 1.34 | 2.5 | $1.7 \cdot 10^{-2}$ | 20288 |
| **Rasl11a** | 22.3 | 0.72 | 1.6 | $1.8 \cdot 10^{-2}$ | 68895 |
| **Npas2** | 2.9 | 0.84 | 1.8 | $1.8 \cdot 10^{-2}$ | 18143 |
| **Actn2** | 18.0 | 0.73 | 1.7 | $1.9 \cdot 10^{-2}$ | 11472 |
| **Mylk4** | 1.5 | 0.84 | 1.8 | $2.0 \cdot 10^{-2}$ | 238564 |
| **Tnni3** | 101.6 | 0.60 | 1.5 | $2.0 \cdot 10^{-2}$ | 21954 |
| **Serpina1b** | 1.8 | 7.84 | 228.4 | $2.1 \cdot 10^{-2}$ | 20701 |
| **Strip2** | 1.7 | 0.85 | 1.8 | $2.2 \cdot 10^{-2}$ | 320609 |
| **Trim63** | 1.2 | 1.60 | 3.0 | $2.3 \cdot 10^{-2}$ | 433766 |
| **Ifit1bl1** | 20.1 | 0.96 | 1.9 | $2.5 \cdot 10^{-2}$ | 667373 |
| **Fndc5** | 9.1 | 0.65 | 1.6 | $2.5 \cdot 10^{-2}$ | 384061 |
| **Thap2** | 5.4 | 0.74 | 1.7 | $2.6 \cdot 10^{-2}$ | 66816 |
| **Ryr2** | 2.4 | 0.63 | 1.6 | $2.8 \cdot 10^{-2}$ | 20191 |
| **Opcml** | 0.4 | 1.09 | 2.1 | $2.9 \cdot 10^{-2}$ | 330908 |
| **Ptpro** | 1.5 | 0.89 | 1.9 | $3.7 \cdot 10^{-2}$ | 19277 |
| **Csrp3** | 23.6 | 0.83 | 1.8 | $3.8 \cdot 10^{-2}$ | 13009 |
| **Gdpd2** | 4.3 | 0.92 | 1.9 | $3.9 \cdot 10^{-2}$ | 71584 |
| **Oas1g** | 4.4 | 0.78 | 1.7 | $4.0 \cdot 10^{-2}$ | 23960 |
| **Klhl41** | 1.4 | 1.22 | 2.3 | $4.0 \cdot 10^{-2}$ | 228003 |
| **Rbm20** | 1.2 | 0.79 | 1.7 | $4.1 \cdot 10^{-2}$ | 73713 |
| **Ccl28** | 0.5 | 1.58 | 3.0 | $4.6 \cdot 10^{-2}$ | 56838 |

**Supplementary Table 3.**
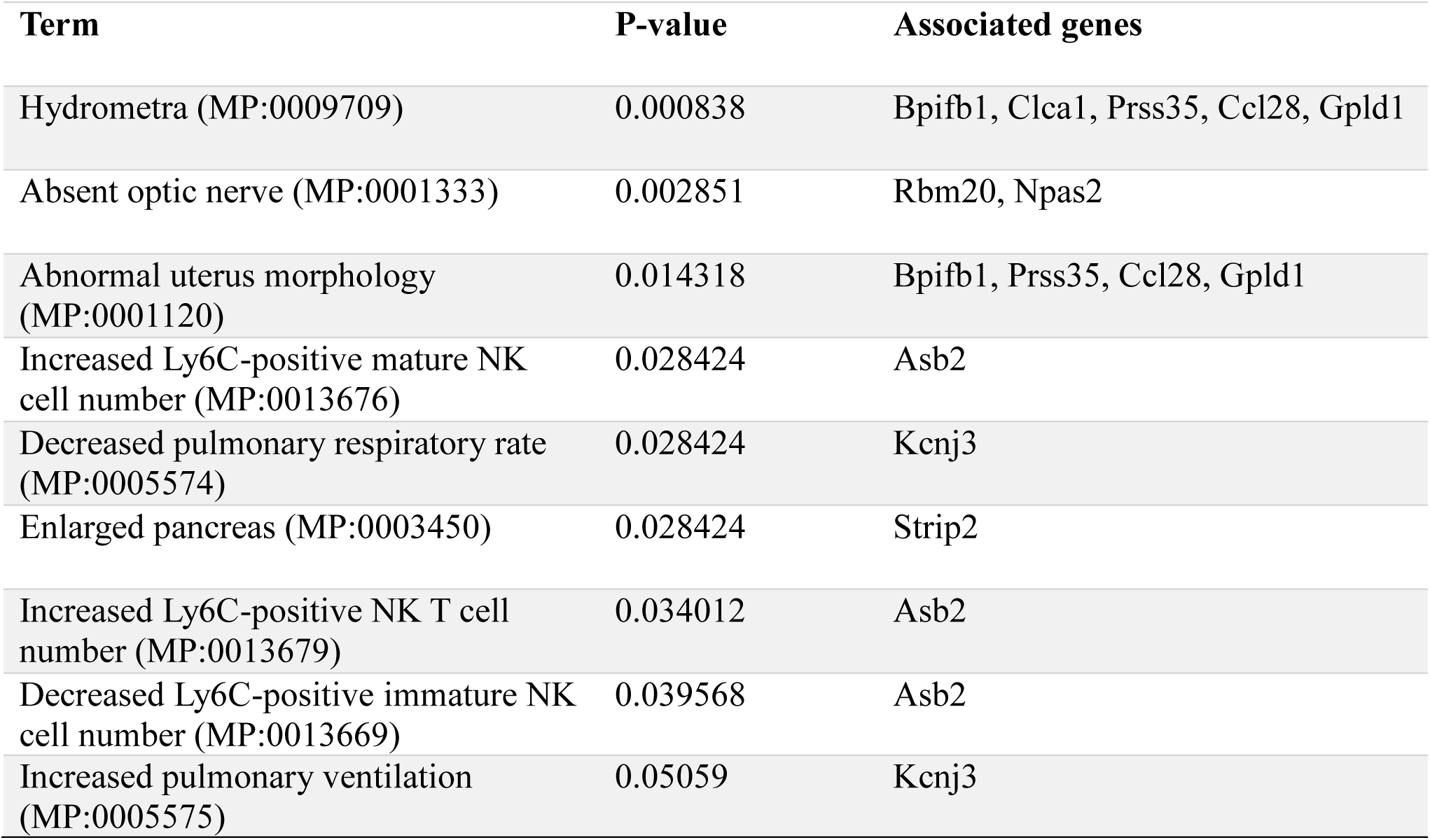
GO analysis of genes that had higher expression in muMT mice compared to WT mice at 4 hpi. Top significant terms are shown based on *P*-values. Analysis was done based on the Knock Out Mouse Phenotyping Program (KOMP2) 2022 gene set library using Enrichr (https://amp.pharm.mssm.edu/Enrichr/).

| Term | P-value | Associated genes |
| --- | --- | --- |
| Hydrometra (MP:0009709) | 0.000838 | Bpifb1, Clca1, Prss35, Ccl28, Gpld1 |
| Absent optic nerve (MP:0001333) | 0.002851 | Rbm20, Npas2 |
| Abnormal uterus morphology (MP:0001120) | 0.014318 | Bpifb1, Prss35, Ccl28, Gpld1 |
| Increased Ly6C-positive mature NK cell number (MP:0013676) | 0.028424 | Asb2 |
| Decreased pulmonary respiratory rate (MP:0005574) | 0.028424 | Kcnj3 |
| Enlarged pancreas (MP:0003450) | 0.028424 | Strip2 |
| Increased Ly6C-positive NK T cell number (MP:0013679) | 0.034012 | Asb2 |
| Decreased Ly6C-positive immature NK cell number (MP:0013669) | 0.039568 | Asb2 |
| Increased pulmonary ventilation (MP:0005575) | 0.05059 | Kcnj3 |

